# Evolution of male costs of copulation in sepsid flies (Diptera: Sepsidae)

**DOI:** 10.1101/265439

**Authors:** Patrick T. Rohner, Kai Shen Yoong, Mindy J. M. Tuan, Rudolf Meier

## Abstract

Reproduction is well known to be costly for females, but longevity costs of copulations in males are still poorly understood. In particular, the effect of the number of copulations on male longevity is rarely considered. Work on black scavenger flies (Diptera: Sepsidae) showed contrasting results: in *Saltella sphondylii* the number of copulations is strongly negatively correlated with male longevity, whereas in *Sepsis cynipsea* mated males did not suffer from reduced longevity. Here we summarize the findings of several studies covering four additional species of sepsid flies from across the phylogenetic tree of sepsids to better understand the evolution of male reproductive costs in this clade. After accounting for the mating system differences between species, we find no evidence for longevity costs in *Allosepsis* sp., *Sepsis fulgens* and *Themira superba*, while in *Saltella nigripes* multiple copulations drastically reduced longevity. Mapping this trait onto the most current phylogenetic hypothesis for Sepsidae suggests that male cost in *Saltella* is derived while there is an absence of longevity costs for all other sepsids. We discuss the origin of this novel longevity cost in *Saltella* in the context of a change in their reproductive strategy, namely the evolution of high polygynandry coupled with unusually brief copulations.

## Introduction

Natural resources are finite and hence investment in one trait often comes at the expense of another. Life history theory predicts that individuals should aim to optimize fitness by trading off energy between traits. Reproduction, being the essential target of direct Darwinian selection, is no exception and can evoke costs in other areas of an organism’s life (Prowse and Partridge 1997; Scharf et al. 2013).

Reproductive costs are well studied in females (Fowler and Partridge 1989; Tatar et al. 1993) with the most obvious one being egg production. Less obvious but also significant are physical injuries suffered during copulation (sepsid flies: Blanckenhorn et al. 2002; eg. traumatic insemination; bean weevils: Crudgington and Siva-Jothy 2000; fruit flies: Kamimura 2007; spiders: Rezac 2009) which may be accompanied by fitness costs caused male seminal fluid (Chapman et al. 1995; Wigby and Chapman 2005) (e.g., transfer of micropathogens (see Poiani 2006)). Other reproductive costs in females include male harassment (Arnqvist 1992; Boness et al. 1995), (postnatal) parental care (Fisher and Blomberg 2011; Visser and Lessells 2001) and the need for morphological adaptations such as elaborate sperm storage organs (Puniamoorthy et al. 2010).

In contrast, male investments in reproduction are less frequently studied. What is known suggests that the male ejaculate is costly because it is not uniform and investment depends on both male and female mating status (Boorman and Parker 1976; Reinhardt 2007) as well as age and fecundity (Lupold et al. 2011). Sperm viability also decreases with male age (Dowling and Simmons 2012). However, reproductive costs of males include many additional factors. For instance, in black scavenger flies (Diptera: Sepsidae), a family of small acalyptrates, most species have developmental costs related to developing elaborate sexual dimorphisms (Ang et al. 2008; Bowsher et al. 2013; Ingram et al. 2008; Rohner et al. 2018). In addition, males need to engage in energy- and time-consuming behavior to obtain copulations. This includes sexual signaling and courtship (eg. Eberhard 2003; Puniamoorthy et al. 2009), perfuming of females with potentially expensive substances (Araujo et al. 2014), competition with other males for mates (eg. Eberhard 2002; Parker 1972), and/or mate guarding (Parker 1972; Ward 1983). Quantifying the costs of these various male investments is difficult, but an overall cost assessment can be obtained by studying the relationship between the number of copulations and male longevity, subsuming the trade-off between reproductive investment and somatic maintenance.

Unfortunately, such data are very difficult to obtain in the field given that the flies are vagile and can live for several weeks. Male copulation costs are thus initially best studied using laboratory experiments. These generally offer better control over factors such as the number of copulations, social context, and intensity of courtship behaviors and can yield hypotheses that can then be tested under field conditions. However, unfortunately even under laboratory conditions it is often difficult to control all desirable factors (e.g., remating rates of males) because not all species have the same mating systems and differences in degree of polygyny and female propensity to copulate influence the ability to carry out identical experiments across species. Yet, including species with different mating systems is desirable in order to test the generality of male costs across whole clades.

### Male costs of reproduction in Sepsidae

One of the first studies demonstrating male longevity costs of reproduction was conducted on *Saltella sphondylii,* a black scavenger fly. The study experimentally controlled the number of copulations in males and then determined male longevity. Martin and Hosken (2004) showed that virgin males lived longer than males engaging in sexual reproduction. By manipulating the number of copulations for each experimental male, *S. sphondylii* virgins lived for about 27.90 (± 1.76) days, whereas males that successfully mated 1, 2, 4 and 6 times suffered from an average lifespan reduction of 3, 6, 7 and 12 days respectively (Martin and Hosken 2004). While this study convincingly demonstrated longevity costs in a sepsid fly, a more recent study on another sepsid, *Sepsis cynipsea,* found no association between the number of copulations and male longevity (Teuschl et al. 2010). These conflicting results within Sepsidae suggest that copulation costs for males can evolve, making sepsids a promising system to study the evolution of male reproductive costs because the evolutionary drivers underlying such variation are not well understood although being crucial for our understanding of mating system evolution (Shuker and Simmons 2014).

We here summarize the results of several studies investigating male longevity costs associated with the number of copulations in four additional species of sepsid flies. The experimental designs between the studies had to accommodate the intrinsic differences in mating systems and behavior between the species. Nevertheless, all studies yielded sufficient information on male mating costs to address the question whether the strong pattern observed in *Saltella* is found throughout Sepsidae. We use the results of these case studies and qualitatively map these data onto a phylogenetic hypothesis of Sepsidae. Given the stark interspecific variation in the elaborateness of copulatory courtship (Puniamoorthy et al. 2009), which can affect male longevity (eg. Papadopoulos et al. 2010), we also test whether variation in courtship intensity mediates longevity costs in males within and across species. In this context, we also present novel data on the courtship behavior of *Saltella nigripes* derived from frame-by-frame video analyses.

## Material and methods

### Taxon sampling

In order to cover the phylogeny as broadly as possible, we studied four species belonging to different major clades of Sepsidae whose relationships have been reconstructed based on egg (Meier 1995b), larval (Meier 1995a), adult (Meier 1996), as well as molecular data for 10 genes (Zhao et al. 2013). The poorly known *Saltella nigripes* Robineau-Desvoidy, 1830 was chosen due to its close relationship to the previously studied *Saltella sphondylii. Themira superba* (Haliday, 1833) represents a major clade that is basal relative to *Saltella* based on morphology and the latest molecular phylogeny, while the species complex *Allosepsis* sp. (Wiedemann, 1824) is more closely related to the largest genus of Sepsidae, *Sepsis* (see fig. 1). Lastly, *Sepsis fulgens* Meigen, 1826 was chosen as a second representative of this species rich genus (fig. 1).

**Figure 1:**
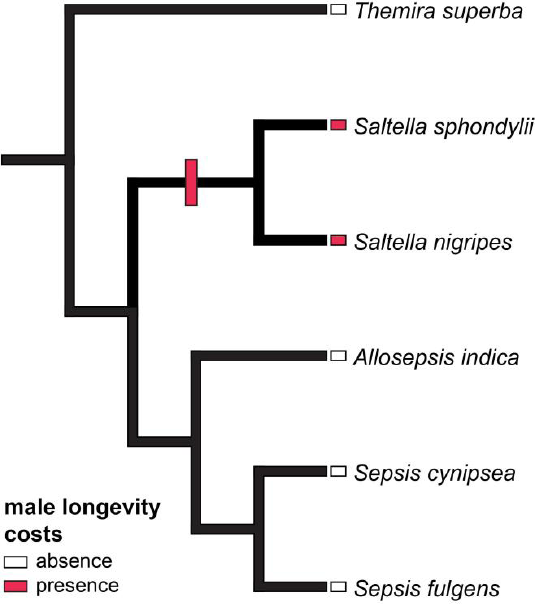
Phylogenetic relationship of the sepsids studied following Zhao et al. (2013). The male reproductive costs observed in both species of *Saltella* represent an autapomorphy of the genus (origin marked in red). These costs are absent in the other studied species marked in white (data on *Saltella sphondylii:* Martin and Hosken 2004; *Sepsis cynipsea:* Teuschl et al. (2010)).

### General laboratory breeding and experimental procedure

Wild-caught females of all species were used to establish laboratory cultures at the National University of Singapore *(Allosepsis, S. fulgens* and *T. superba)* and the University of Zurich *(Saltella nigripes).* All species were identified using the digital reference collection SepsidNet (Ang et al. 2013; Meier 2017). Cultures were housed for several generations in plastic containers equipped with sugar, water and breeding substrate *ad libitum.* To fulfill the ecological requirements of each species, we kept *Allosepsis* sp. (Indonesia strain) and *S. fulgens* (Sweden) at 28.0°C while *T. superba* (Germany) and *Saltella nigripes* (Switzerland) were kept at 20.0°C. In order to obtain virgin flies for experimental manipulation, breeding substrate was provided to all parental cultures *(Allosepsis* sp., *S. fulgens, Saltella nigripes:* cow dung; *T. superba*: waterfowl-cow dung mix). Dung was replaced every 24 hours over a span of several days, providing a continuous supply of virgin flies for mating trials. To ensure virginity, emerged flies were separated by sex within 24 hours. To eliminate the effect of intra-sexual interactions on longevity, males were kept individually in small vials, equipped with water, sugar and dung. Females were kept in groups in larger containers. As a control for the effect of any experimental treatment, virgin males that had no exposure to female flies throughout their life were monitored in separate cages.

Variation in the number of times a male copulates is intrinsically confounded by the presence of a female, but controlling for the latter is difficult. In several sepsids, non-fertile females show reluctance behavior towards mating and can repel mates successfully. Such reluctant females are thus well suited to account for female presence. We successfully applied this procedure in *Allosepsis*, while this manipulation was not feasible in the other species. In order to receive reluctant *Allosepsis* females, some virgins were kept without dung as females without access to dung do not develop eggs and are unlikely to mate (Teuschl and Blanckenhorn 2007), showing high levels of reluctance behavior.

### Mating Trials

Mating trials were conducted once males were sexually mature (4 to 5 days old). During each mating trial, males were removed from their individual chambers and placed with females of known age into a small vial (with the exception of *Saltella nigripes* for which not enough females were available, all females were virgin). Due to intrinsic differences in mating behavior and the mating system, experimental protocols had to be customized for each species as outlined in the following.

### Allosepsis *sp*

Sexually mature virgin females usually mate within 10 minutes of being exposed to any male. Thus, males could be randomly assigned to 0, 1, 2, or 3 copulation treatments prior to the start of experiments. During each mating trial, males were provided with a 4-day old virgin female for 4 hours. Mating usually concluded within 30 minutes of contact. If no copulation occurred within 30 minutes, the female was replaced with another virgin female. To control for the presence of a female, males were provided with either a fertile or a non-fertile female (control treatment). When exposed to non-fertile females, males still courted but were rejected by females (and mating rate dropped to less than 20%). Males that did not copulate according to the assigned treatment group were discarded (10 males throughout the experiment). After mating trials (or upon emergence for control males), males were checked every 12 hours for survival. The time of death was recorded to estimate the longevity of each experimental male. Body size (scutum width) for each experimental male was measured to account for its potentially confounding effects on longevity.

### Sepsis fulgens

This generalist species is very common in Central Europe and occurs to the thousands near pig manure or dung heaps. Although densities are high, copulations are only rarely observed (pers. obs. PTR) which is also reflected in rather low virgin mating success under laboratory conditions (<35%). Therefore, it was not possible to assign treatment groups *a priori.* We thus allowed males to mate freely and assigned treatment groups after all mating trials were performed according to the number of successful copulations. Mate trials were conducted when the males were 5, 7, 9, 11 and 13 days old in order to have a sufficiently large number of individuals with different copulation numbers. Each trial lasted for about 90 minutes and was recorded using a Canon Legria HF S30 video camera. To maximize copulation rates, males were provided with two 6-day old virgin females. Copulation normally occurred within 45 minutes. Male mating effort was estimated using two measures: (1) the number of times a male mounted on the female but was shaken off and (2) copulation duration. A total of 118 trials, each of ca. 90 minutes duration, were conducted to quantify copulation duration in seconds and number of time the males tried to mount the females. After mating trials (or upon emergence for control males), males were checked every 24 hours for survival. The time of death was recorded to estimate the longevity of each experimental male. Body size (scutum width) for each experimental male was measured to account for its potentially confounding effects on longevity

### Themira superba

Similar to some other species of the genus *Themira, T. superba* prefers waterfowl dung as breeding substrate and its distribution is thus rather spotty and often found near lakes, river shores or other dull places. Similar to *Allosepsis,* this species has a high virgin mating rate (> 90%), however, due to skewed sex ratios in the laboratory populations, male copulations cannot be controlled as well as in *Allosepsis*, and the treatments were assigned after mating trials, similarly to *S. fulgens.* Each male was presented with two virgin females during mating trials that lasted for two hours. After mating trials, males were checked every 12 hours for survival. The time of death was recorded to estimate the longevity of each experimental male. Body size (scutum width) for each experimental male was measured to account for its potentially confounding effects on longevity.

### Saltella nigripes

This is a rather large, dull-looking sepsid fly associated to cow dung in Central Europe. This species is generally rare in the wild throughout the season but sporadically occurs at high densities near cow dung in autumn. During such occasions, pairs can be observed mating at high frequencies near dung pads whereas both sexes remate readily (pers. obs. PTR), which is a highly unusual behavior (c.f. Pont & Meier, 2002). This species is difficult to breed under laboratory conditions due to high larval and adult mortality. We were thus forced to modify the experimental protocol to fit these constraints (although treatments were identical unless stated otherwise). Males were *a priori* split into a control group, which never had contact with females, and a treatment group in which single males were paired with a single female. Mating trials were conducted twice daily over the course of five days, always resulting in ten successful copulations per male in total. Although this may seem a particularly high number of matings, field observations suggest that this highly polygynandrous species indeed copulates very frequently. Due to the low number of available females, not all females were virgin, but due to the high mating rates virgin matings are likely to be the exception in the field. Related to this species’ unusual mating system, reluctant females could not be obtained and hence controlling for the presence of females was not feasible.

Following the experimental procedure described in Puniamoorthy et al. (2008) (see also: Puniamoorthy et al. 2009; Tan *et al.* 2011), we further recorded the mating behavior of *Saltella nigripes* using a Nikon DS-Fi1 digital video camera (n = 11 copulations). Mating behavior was described using the character definition of Puniamoorthy et al. (2009a) and its extension by Tan et al. (2011).

### Statistical analysis

Cox Proportional Hazard models were used to test for differences in survival between treatment groups. Four variables were included in the analysis: treatment (number of copulations: 0 – 10), exposure to females (yes or no), batch number (1-3; blocking effect), and body size. Helmert contrasts were specified to detect differences between treatment groups (see table 1). For *S. fulgens,* two additional variables were included: duration of copulation, and number of mountings were added to the analysis. All analyses were conducted using SPSS v22.

**Table 1:**
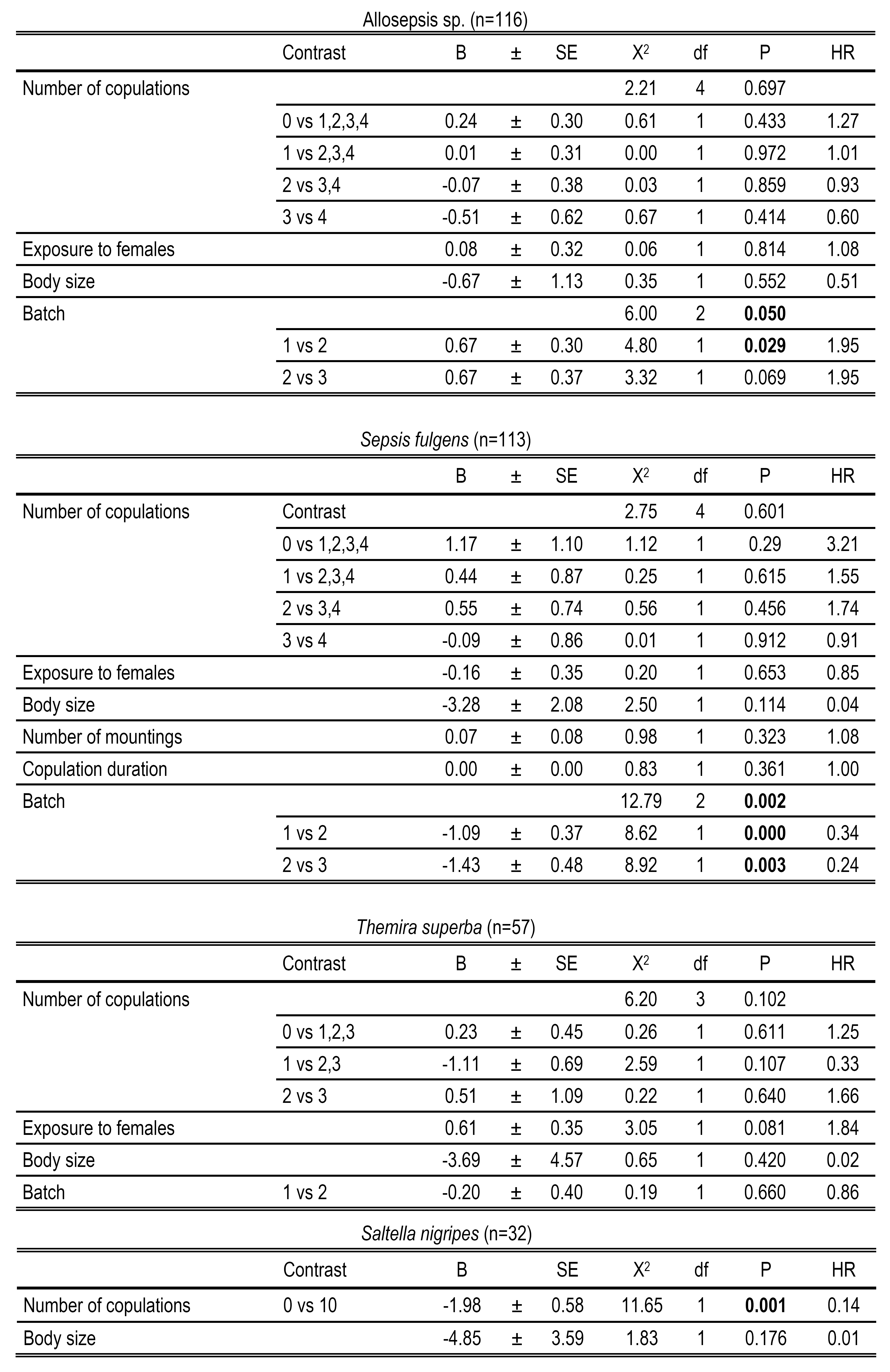
Results of the Cox Proportional Hazard model using male longevity as dependent variable (HR: hazard ratio).

| *Allosepsis* sp. (n=116) |  |  |  |  |  |  |  |  |
| --- | --- | --- | --- | --- | --- | --- | --- | --- |
|  | Contrast | B | ± | SE | X<sup>2</sup> | df | P | HR |
| Number of copulations |  |  |  |  | 2.21 | 4 | 0.697 |  |
|  | 0 vs 1,2,3,4 | 0.24 | ± | 0.30 | 0.61 | 1 | 0.433 | 1.27 |
|  | 1 vs 2,3,4 | 0.01 | ± | 0.31 | 0.00 | 1 | 0.972 | 1.01 |
|  | 2 vs 3,4 | -0.07 | ± | 0.38 | 0.03 | 1 | 0.859 | 0.93 |
|  | 3 vs 4 | -0.51 | ± | 0.62 | 0.67 | 1 | 0.414 | 0.60 |
| Exposure to females |  | 0.08 | ± | 0.32 | 0.06 | 1 | 0.814 | 1.08 |
| Body size |  | -0.67 | ± | 1.13 | 0.35 | 1 | 0.552 | 0.51 |
| Batch |  |  |  |  | 6.00 | 2 | **0.050** |  |
|  | 1 vs 2 | 0.67 | ± | 0.30 | 4.80 | 1 | **0.029** | 1.95 |
|  | 2 vs 3 | 0.67 | ± | 0.37 | 3.32 | 1 | 0.069 | 1.95 |
| *Sepsis fulgens* (n=113) |  |  |  |  |  |  |  |  |
|  | Contrast | B | ± | SE | X<sup>2</sup> | df | P | HR |
| Number of copulations |  |  |  |  | 2.75 | 4 | 0.601 |  |
|  | 0 vs 1,2,3,4 | 1.17 | ± | 1.10 | 1.12 | 1 | 0.29 | 3.21 |
|  | 1 vs 2,3,4 | 0.44 | ± | 0.87 | 0.25 | 1 | 0.615 | 1.55 |
|  | 2 vs 3,4 | 0.55 | ± | 0.74 | 0.56 | 1 | 0.456 | 1.74 |
|  | 3 vs 4 | -0.09 | ± | 0.86 | 0.01 | 1 | 0.912 | 0.91 |
| Exposure to females |  | -0.16 | ± | 0.35 | 0.20 | 1 | 0.653 | 0.85 |
| Body size |  | -3.28 | ± | 2.08 | 2.50 | 1 | 0.114 | 0.04 |
| Number of mountings |  | 0.07 | ± | 0.08 | 0.98 | 1 | 0.323 | 1.08 |
| Copulation duration |  | 0.00 | ± | 0.00 | 0.83 | 1 | 0.361 | 1.00 |
| Batch |  |  |  |  | 12.79 | 2 | **0.002** |  |
|  | 1 vs 2 | -1.09 | ± | 0.37 | 8.62 | 1 | **0.000** | 0.34 |
|  | 2 vs 3 | -1.43 | ± | 0.48 | 8.92 | 1 | **0.003** | 0.24 |
| *Themira superba* (n=57) |  |  |  |  |  |  |  |  |
|  | Contrast | B | ± | SE | X<sup>2</sup> | df | P | HR |
| Number of copulations |  |  |  |  | 6.20 | 3 | 0.102 |  |
|  | 0 vs 1,2,3 | 0.23 | ± | 0.45 | 0.26 | 1 | 0.611 | 1.25 |
|  | 1 vs 2,3 | -1.11 | ± | 0.69 | 2.59 | 1 | 0.107 | 0.33 |
|  | 2 vs 3 | 0.51 | ± | 1.09 | 0.22 | 1 | 0.640 | 1.66 |
| Exposure to females |  | 0.61 | ± | 0.35 | 3.05 | 1 | 0.081 | 1.84 |
| Body size |  | -3.69 | ± | 4.57 | 0.65 | 1 | 0.420 | 0.02 |
| Batch | 1 vs 2 | -0.20 | ± | 0.40 | 0.19 | 1 | 0.660 | 0.86 |

*Saltella nigripes* (n=32)
|  | Contrast | B |  | SE | X<sup>2</sup> | df | P | HR |
| --- | --- | --- | --- | --- | --- | --- | --- | --- |
| Number of copulations | 0 vs 10 | -1.98 | ± | 0.58 | 11.65 | 1 | **0.001** | 0.14 |
| Body size |  | -4.85 | ± | 3.59 | 1.83 | 1 | 0.176 | 0.01 |

## Results

### Male costs of copulations

Cox Proportional Hazard models revealed no impact of the number of copulations on longevity in *Allosepsis* sp., *Sepsis fulgens* and *Themira superba* (table 1. fig. 2). None of the other predictor variables had a significant effect on survival, except for (probably random) differences between batches in *Allosepsis* sp. and *Sepsis fulgens* (table 1). Also, male mating effort, estimated by copulation duration and the number of unsuccessful mountings, had no impact on survival in *Sepsis fulgens* (table 1).

In sharp contrast, *Saltella nigripes* males which copulated with females suffered from a pronounced reduction of lifespan (longevity: virgin: 30.1 +/-7.6 days; mated: 18.8 +/-0.1 days; table 1, fig. 2). We note that we could not control for the presence of females due to the intrinsically high remating frequency of females and we can thus not disentangle its effect from the cost of copulation *per se.* Also, with ten copulations per male (although mimicking natural remating rates) data are not directly comparable to other species in terms of physiological cost per unit mating (see Discussion).

**Figure 2:**
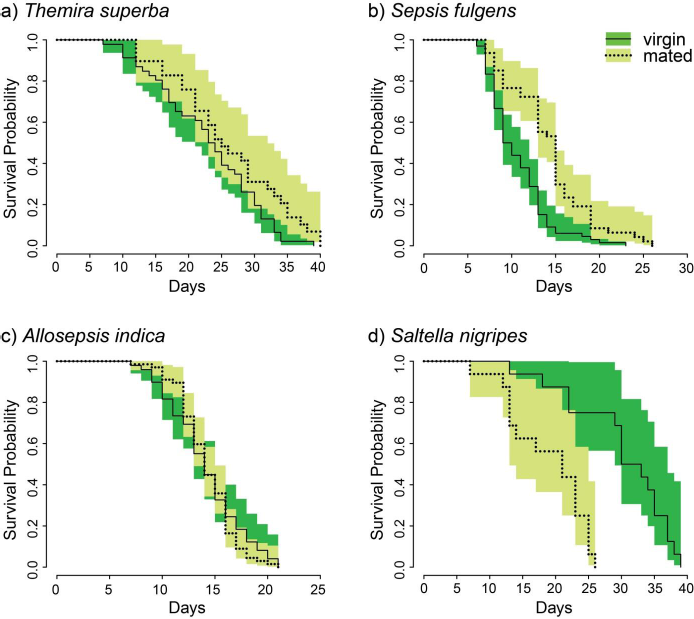
Survival probability for four closely related species of sepsid flies. Colors indicate 95% CI separated by mating status of males (light green: virgins; dark green: mated). *Themira superba* (a), *Sepsis fulgens* (b) and *Allosepsis* sp. (c) do not suffer from reduced viability after mating. In contrast, virgin *Saltella nigripes* live much longer than mated conspecifics (d).

### Courtship behavior of Saltella nigripes

Frame-by-frame analysis of video recordings revealed little intraspecific variation in copulatory courtship (as is common in sepsids, Puniamoorthy et al. 2009). As in *Saltella sphondylii,* copulation duration in *S. nigripes* is unusually short (3-4 minutes). The elaborate pre-copulatory courtship known from other sepsids is absent in both *Saltella* species. Males climb onto the female and usually engage in copulations immediately. However, in contrast to *Saltella sphondylii,* copulations are not characterized by multiple insertions of the aedeagus (i.e. copulations) in *S.nigripes:* even though genital contact is terminated after 3-4 minutes, the male will not dismount the female until after a period of 8 to 15 minutes of post-copulatory courtship. Moreover, the courtship repertoire differs strongly between the two closely related species: *Saltella nigripes* does not use proboscis stimulation of females (character 4; cf. Tan *et al.,* 2011), midlegs are frequently stretched away from the female during copulations (i.e. ‘balancing’, character 7; cf. Tan *et al.,* 2011), and the surstyli are not used to stimulate the female abdomen (character 25; cf. Tan *et al.,* 2011). As in *sphondylii,* male *nigripes* rub their midlegs against their hindlegs and proceed with rubbing the female’s head with their midtarsi; however, this movement is performed in an alternate fashion whereas in *sphondylii* both midlegs are used to rub the female’s head simultaneously (character 9; cf. Tan *et al.,* 2011).

The females also differ in their copulatory behavior: female shaking was never observed in *S. nigripes* (character 28; cf. Tan *et al.,* 2011). Females were never observed ‘kicking’ the male, and opposite to *sphondylii* (character 31; cf. Tan *et al.,* 2011), which never eject their ovipositor directly prior or after copulations, all female *S. nigripes* ejected their ovipositor directly after genital contact (character 32; cf. Tan *et al.,* 2011). Synapomorphies of *Saltella spp.* comprise the quick separation after copulation, which does not involve turns by the male (character 27; cf. Tan *et al.,* 2011), as well as a tandem walk during copulation (character 19; cf. Tan *et al.,* 2011).

## Discussion

In sepsid flies, mating systems vary strongly between species and sometimes even among populations (Puniamoorthy et al. 2012; Rohner et al. 2016) with great variation in their elaborate copulatory courtship displays (Puniamoorthy et al. 2008, 2009). Sepsids are nevertheless occasionally cited as a classic example for male longevity costs associated with multiple copulations although this perception is based on a single species, *Saltella sphondylii* (Martin and Hosken 2004) whose mating system is exceptional in many ways (Tan et al., 2010). We here find no evidence for drastic longevity costs in other sepsid species representing those clades that include more than 90% of the species diversity *(Allosepsis, T. superba, S. fulgens* (this study) and *S. cynipsea* studied in Teuschl et al. (2010)) but find such mating costs in *Saltella nigripes,* a close relative of *S. sphondylii.* It thus appears likely that the live-fast-die-young life history of *Saltella* is an exception and restricted to a species-poor clade within Sepsidae (fig.1).

Our conclusion of “no evidence for drastic longevity costs” is however to be taken with some caution because the mating behavior of the studied species differed by so much that it proved impossible to apply the exact same experimental conditions. With regard to *Saltella nigripes*, we had to use non-virgin females for testing and used a relatively high number of copulations which mimics the polygyny observed in the wild. In *Sepsis fulgens* and *Themira superba*, female resistance is unpredictable and often very high, which rendered it exceedingly difficult to assign males a priori to different treatment classes (= number of copulations). Instead, we evaluated whether there was a copulation cost to males with a larger number of copulations. This means that we cannot rule out that weak males avoided multiple copulations; i.e., damage or expenditure caused by previous copulations led weak males to drop out of the mating game and/or costs would have become apparent if they had been forced to copulate multiple times. Note however, that our conclusion that there is no male copulation cost for *S. fulgens* is also consistent with the study of *S. cynipsea* by Teuschl et al. (2010) who strictly controlled for the number of copulations and yet could also not reject the null hypothesis of no male longevity cost. The lack of effect in *S. fulgens* and *T. superba* is thus likely to reflect an actual lack of physiological cost and not a consequence of male bet hedging.

Overall, we find evidence for male copulation costs in *Saltella* but not the other genera, offering an opportunity to investigate the underlying ultimate and proximate drivers of such variation. Males suffer from viability costs mediated by reproductive investment in a large array of species (Harwood et al. 2015; Scharf et al. 2013) but even among closely related species, male longevity costs can vary in its extent and may not always be present (eg. in ladybird beetles; see Perry and Tse (2013) and references therein). In sepsids, evidence for male longevity costs exists for *Saltella sphondylii* (Martin and Hosken 2004) but is absent in all other studied sepsids with the exception of its congener *Saltella nigripes* (this study). Character optimization on the latest phylogenetic hypothesis for Sepsidae hence suggests that the longevity costs in *Saltella* are a derived trait. Based on observations in the laboratory, the high levels of polyandry in *Saltella* are very unlikely to be realized in any species of *Sepsis, Themira* or *Allosepsis.* Given this lack of high remating rates and the lack of copulatory costs under laboratory conditions, copulations should not evoke viability costs under natural conditions and hence may not be of great evolutionary relevance. In contrast, given our field observations, both species of *Saltella* are very likely to copulate ten times or more throughout their lifetime. Longevity costs are thus expected to also arise in the field. The high male mating rates and the considerable male mating costs are therefore likely to be derived and restricted to a species-poor clade including *Saltella* (<10 species) while there is no evidence suggesting the presence of pronounced male mating costs in the vast majority of sepsid species (>300 species).

Interestingly, the mating system of *Saltella* spp. does not only feature an unusually high level of polygynandry, but also the shortest copulation times observed in sepsids (Tan et al., 2010; this study), a lack of reluctance behavior by females (Martin and Hosken 2004; Schulz 1999), and post-copulatory guarding. The latter, which is often linked to the avoidance of sperm competition (Simmons 2014), has been described as an autapomorphy for *S. sphondylii* (Schulz 1999) but is also apparent in *S. nigripes,* where males stay mounted after terminating genital contact and continue with (post-copulatory) courtship. Being rare or entirely absent in all other sepsids studied to date, these traits suggest a pronounced role of sperm competition and/or cryptic female choice in *Saltella.* Post-copulatory selective forces could thus constitute the major ultimate evolutionary drivers for high reproductive investments, outweighing longevity costs in males via high short-term fertilization success. However, this association requires further testing. In addition to high investment into sperm or seminal fluids, longevity costs could also be related to injuries inflicted during copulation, again a mechanism that requires further scrutiny.

### Is courtship costly for male sepsids?

Earlier work on sepsids has revealed great diversity and the fast evolution of behavioral courtship elements (Puniamoorthy 2014; Puniamoorthy et al. 2009; Tan et al. 2010). Such elaborate courtship has been found to be costly in other taxa (e.g. spiders: Hoefler (2008); medflies: Papadopoulos et al. (2010); mosquitoes: South et al. (2009)), but it remains unknown whether male courtship can proximately evoke costs in sepsids. Our behavioral assay on *Saltella nigripes* confirms that courtship behavior must evolve quickly because it differs greatly between the two closely related *Saltella* species. But we cannot identify specific courtship elements in *Saltella* that are likely to account for the interspecific variation in male longevity costs. The abrupt dismount of the male after (the final) copulation as well as the male walking in tandem with the female are observed in both *Saltella* spp. and are absent in the rest of Sepsidae, yet do not give the appearance of being particularly costly. In addition, intraspecific variation in courtship intensity, which we here assessed in detail for *S. fulgens,* was not associated to longevity. Even when adding copulation duration in the analysis of *S. fulgens,* we found no reduction in lifespan. Thus, there is currently no evidence that courtship intensity or mating duration affect male longevity in sepsids. The proximate mechanisms leading to variation in male longevity costs in sepsids therefore remain elusive.

### Conclusions

Comparing our results for *Allosepsis* to previously published data on *Saltella sphondylii* and *Sepsis cynipsea,* we find evidence for interspecific variation in male copulatory costs: While *S. sphondylii* suffers from increased mortality if mated four times (Martin and Hosken 2004), *Sepsis cynipsea* (Teuschl et al. 2010) and *Allosepsis* sp. (this study) do not show reduced longevity when mated equally often. Trying to generalize these results, and tracking its evolution across the phylogeny, we aimed at estimating copulatory longevity costs in three other sepsid species. However, due to the great variation in mating systems among sepsids, we were not able to assign treatments *a priori* for *S. fulgens* and *T. superba*. Viability costs in these species either are thus not pronounced or remain hidden if males only mate if they can afford to mate. The latter scenario would suggest that these species circumvent copulatory costs by behavioral means, again contrasting to the two *Saltella* species, which mate regardless of longevity costs (males engage in copulations almost immediately). Irrespective of the mechanisms preventing or imposing copulatory costs, all available evidence suggests that the male mating costs observed for *Saltella* are an exception. Thus, under the most parsimonious scenario, the pronounced and often cited male longevity costs to copulation observed in the *Saltella* clade (<10 species) are likely derived and not present in the rest of the family, nor in the common ancestor of Sepsidae. The extensive copulatory courtship observed in sepsids is unlikely to carry a high cost for males, but post-copulatory selective agents could promote the evolution of high copulatory investments in male *Saltella.* Future research should aim at identifying the proximate drivers mediating the evolution of male reproductive costs and further broaden the taxon sampling including more species and clades to test our predictions.

## Acknowledgements

We thank the Evolutionary Biology Laboratory at the National University of Singapore and the Invertebrate Behavioral and Evolutionary Ecology group at the University of Zurich for their input and support of this project. RM would like to acknowledge support from MOE Grant R-154-000-476-112.

